# Development of the Incompatible Insect Technique targeting *Aedes albopictus*: introgression of a wild nuclear background restores the performance of males artificially infected with *Wolbachia*

**DOI:** 10.1101/2024.11.25.625297

**Authors:** Quentin Lejarre, Sarah Scussel, Jérémy Esnault, Benjamin Gaudillat, Marianne Duployer, Patrick Mavingui, Pablo Tortosa, Julien Cattel

## Abstract

The bacterium *Wolbachia pipientis* is increasingly studied for its potential use in controlling insect vectors or pests due to its ability to induce Cytoplasmic Incompatibility (CI). CI can be exploited by establishing an opportunistic *Wolbachia* infection in a targeted insect species through trans-infection and then releasing the infected males into the environment as sterilizing agents. Several host life history traits (LHT) have been reported to be negatively affected by artificial *Wolbachia* infection. *Wolbachia* is often considered as the causative agent of these detrimental effects, and the importance of the host’s genetic origins in the outcome of trans-infection is generally overlooked. In this study, we investigated the impact of host genetic background using an *Aedes albopictus* line recently trans-infected with *w*Pip from *Culex pipiens* mosquito, which exhibited some fitness costs. We measured several LHTs including fecundity, egg hatch rate and male mating competitiveness in the incompatible line after four rounds of introgression aiming at restoring genetic diversity in the nuclear genome. Our results show that introgression with a wild genetic background restored most fitness traits and conferred mating competitiveness comparable to that of wild males. Finally, we show that introgression leads to faster and stronger population suppression under laboratory conditions. Overall, our data support that host genome plays a decisive role in determining the fitness of *Wolbachia* infected incompatible males.

**Importance:** The bacterium *Wolbachia pipientis* is increasingly used to control insect vectors and pests through the Incompatible Insect Technique (IIT) inducing a form of conditional sterility when a *Wolbachia*-infected male mates with an uninfected or differently infected female. *Wolbachia* artificial trans-infection has been repeatedly reported to affect mosquitoes LHTs, which may in turn compromise the efficiency of IIT. Using a tiger mosquito (*Aedes albopictus*) line recently trans-infected with a *Wolbachia* strain from *Culex pipiens* and displaying reduced fitness, we show that restoring genetic diversity through introgression significantly mitigated the fitness costs associated with *Wolbachia* trans-infection. This was further demonstrated through experimental population suppression, showing that introgression is required to achieve mosquito population suppression under laboratory conditions. These findings are significant for the implementation of IIT programs, as an increase in female fecundity and male performance improves mass rearing productivity as well as the sterilizing capacity of released males.

## Introduction

The α-proteobacterium *Wolbachia* is among the most abundant symbionts, infecting around 50% of all arthropod species (1–4). This maternally transmitted bacterium induces a wide range of phenotypes in its hosts, including reproductive manipulations to favor its spread in natural populations (5). The most common manipulation induced by *Wolbachia* infection is a form of conditional sterility named Cytoplasmic Incompatibility (CI) (6, 7), which is an embryonic lethality resulting from mating between a *Wolbachia*-infected male and a female that is either uninfected or infected with a distinct incompatible *Wolbachia* strain (8–10). This natural phenomenon can be exploited in biological control though two distinct strategies. *Wolbachia-*infected mosquitoes can be used to replace the existing mosquito population with one less likely to transmit the pathogen, generating long-term reductions in transmission (a *Wolbachia*-based population replacement approach). Alternatively, CI can be exploited to suppress the existing mosquito population by releasing males only, a *Wolbachia*-based population suppression approach also called Incompatible Insect Technique (IIT) (11, 12), in which incompatible males are used as a sterilizing agent to control the population size of the target species (13, 14).

The main advantage of IIT, when compared for example to other techniques such as the sterile insect technique, is that males are “ready to use” and do not require specialized equipment for sterilization. However, IIT is highly sensitive to the accidental release of females, which could lead to the undesirable establishment of *Wolbachia* transinfected mosquitoes. This can be prevented by combining IIT with irradiation, which will sterilize accidentally released females (15, 16), or by running IIT as a standalone (17–19) provided a highly effective sexing system based on genetic (20–23) or mechanical/automated processes (17, 24) is available. Whichever method is employed, the use of a transinfected line exhibiting bidirectional CI with the resident population will drastically reduce the probability of invasion (25).

While initially limited to species where natural *Wolbachia* infections allowed the expression of CI (14, 26–28), or to closely related species allowing horizontal *Wolbachia* transfer through introgression (29), IIT can be considered in a wider range of species through the artificial transfer of CI-inducing *Wolbachia* strains (30–33). *Wolbachia* trans-infected lines have been repeatedly reported to display reduced fitness including reduced fecundity, lower egg hatch rates and/ or survival (34–38). Such fitness costs have been generally attributed to the artificial *Wolbachia* infection. It has been proposed that these costs could be attenuated by artificial selection expected to stabilize the density of the symbiont together with its distribution in the tissues of the new host (39–42). Alternatively, it has also been proposed that the bottleneck in host diversity induced by the trans-infection process may be responsible for some of the measured deleterious effects, although this has been rarely tested to date (32, 43). In most cases, the selection of one (or few) isofemale lines showing higher maternal transmission during the first generations following trans-infection will result in a loss of genetic population diversity.

In this study, we addressed the importance of host genetic background on the performance of *Wolbachia* trans-infected lines by comparing different LHTs in an *Aedes albopictus* trans-infected line to the same line following a few rounds of introgression, or to the wild type line used for this introgression. Finally, we compared the capacity of incompatible males to suppress a population in laboratory-controlled conditions. The data presented show that a small number of rounds of introgression are sufficient to suppress fitness costs associated with *Wolbachia* trans-infection.

## Results

### CI penetrance of wPip-IV in an introgressed background

The mosquito lines used in the experiments include (i) a doubly *w*AlbA/B infected line, referred herein as wild-*w*Alb, (ii) the *w*Pip trans-infected line that was either introgressed with wild-*w*Alb and referred as wild-*w*Pip, or (iii) not introgressed and referred as lab-*w*Pip. Reciprocal crosses between wild-*w*Alb and wild-*w*Pip lines confirmed that *w*Pip-IV induced bidirectional CI in *Ae*. *albopictus*. Indeed, when crossed with wild-*w*Alb females, males from wild-*w*Pip induced complete CI (0% egg hatching rate). The reciprocal crosses between males from wild-*w*Alb line and females from wild-*w*Pip displayed an average hatch rate of 0.46% (±0.46) (Table 1). The same pattern was observed when using the non introgressed line (20), showing, as expected, that introgression did not modify CI.

**Table 1.** Hatch rate obtained in control and reciprocal crosses between the transinfected wild-*w*Pip and the wild-*w*Alb line. The number of eggs counted per cross is indicated in brackets.

| crosses | hatch rate (%) (±SD) | CI (%) | CIcorr (%) |
| --- | --- | --- | --- |
| ♂ *wild-wPip* x ♀ *wild-wAlb* | 0.00 ($\pm 0.00$) (9825) | 100 | 100 |
| ♂ *wild-wAlb* x ♀ *wild-wPip* | 0.46 ($\pm 0.46$) (3865) | 99.54 | 99.49 |
| ♂ *wild-wAlb* x ♀ *wild-wAlb* | 91.22 ($\pm 0.20$) (3447) | | |
| ♂ *wild-wPip* x ♀ *wild-wPip* | 89.86 ($\pm 0.04$) (3512) | | |

### Effect of a wild genetic background introgression on life history traits (LHT)

We assessed the effect of introgression on the LHTs of the trans-infected line that was either introgressed or not introgressed with the wild-*w*Alb line. LHTs included longevity, number of laid eggs, hatching rate and the insemination male capacity. For both sexes, the lab-*w*Pip showed the lowest longevity with a significant difference compared to the two other wild-*w*Pip and wild-*w*Alb lines (*P*<0.001 for each comparison, Fig. 1A and 1B), for which a similar longevity was observed. We also observed a significant difference in the hatch rates between the lab-*w*Pip, displaying an average hatch rate of 56.03% (±28.83) and both wild-*w*Pip and wild-*w*Alb lines, for which similar hatch rates were observed, 78.88% (±23.58) and 82.77% (±18.04) respectively (*P*=0.681) (Fig. 1C). The number of eggs laid per female was not significantly different between lines although higher for females from the wild-*w*Alb line that averaged 70.55 (±30.24) eggs per female, compared to 62.64 (±28.86) and 61.65 (±25.53) for females from the wild-*w*Pip and lab-*w*Pip lines, respectively (Fig. 1D). The insemination capacity of males was evaluated by crossing males and females from the same line and by crossing incompatible males from the wild-*w*Pip or lab-*w*Pip with wild-*w*Alb females. Of note, over 99,5% of dissected inseminated females (957/960) had two filled spermathecae, aligning with a pattern typically reported for *Aedes albopictus* (44, 45). As expected, decreasing the female/male ratio reduced the percentage of inseminated females for all three mosquito lines regardless of the combination of males and females (Fig. 2A & B). For all 3 tested ratios, males from wild-*w*Pip and wild-*w*Alb lines presented a similar insemination capacity when crossed with females from their own line (Fig. 2A). Interestingly, we also observed a similar insemination capacity between these males when crossed with wild-*w*Alb females (Fig. 2B). Finally, the insemination capacity of lab-*w*Pip males was lower than that of males from a wild genetic background (wild-*w*Alb and wild-*w*Pip).

**FIG 1.**
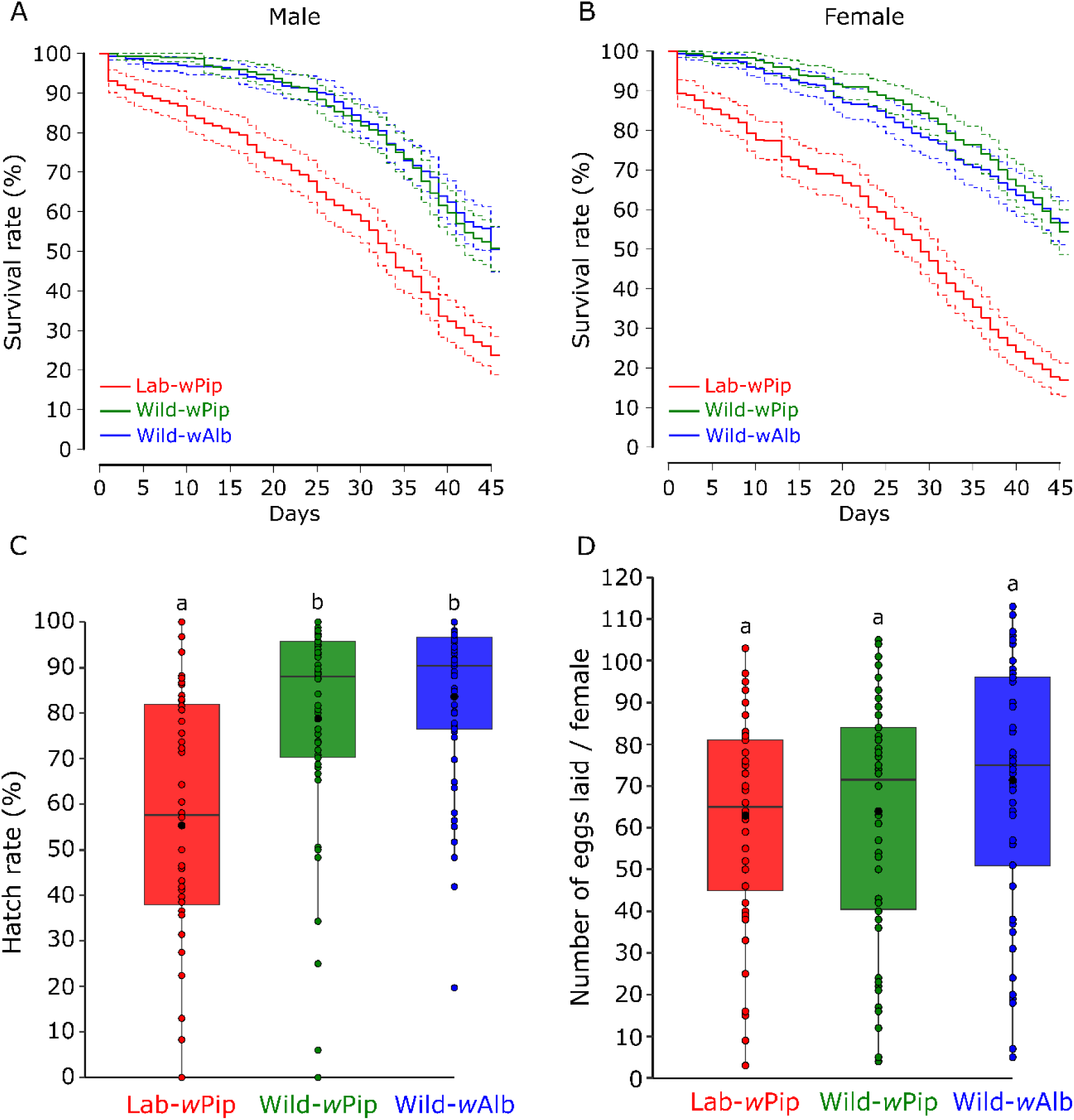
Life history traits of trans-infected lab-*w*Pip and wild-*w*Pip lines compared to wild-*w*Alb line. A: Male longevity. B: Female longevity. C: Eggs hatching rate (%). D: Number of eggs laid per female. For FIG1A & 1B, the solid lines indicate the mean values and the dotted lines the 95% confidence interval. For FIG 1C & 1D, black dots indicate mean values, horizontal bars the medians, and distinct letters significant variations between lines.

**FIG 2.**
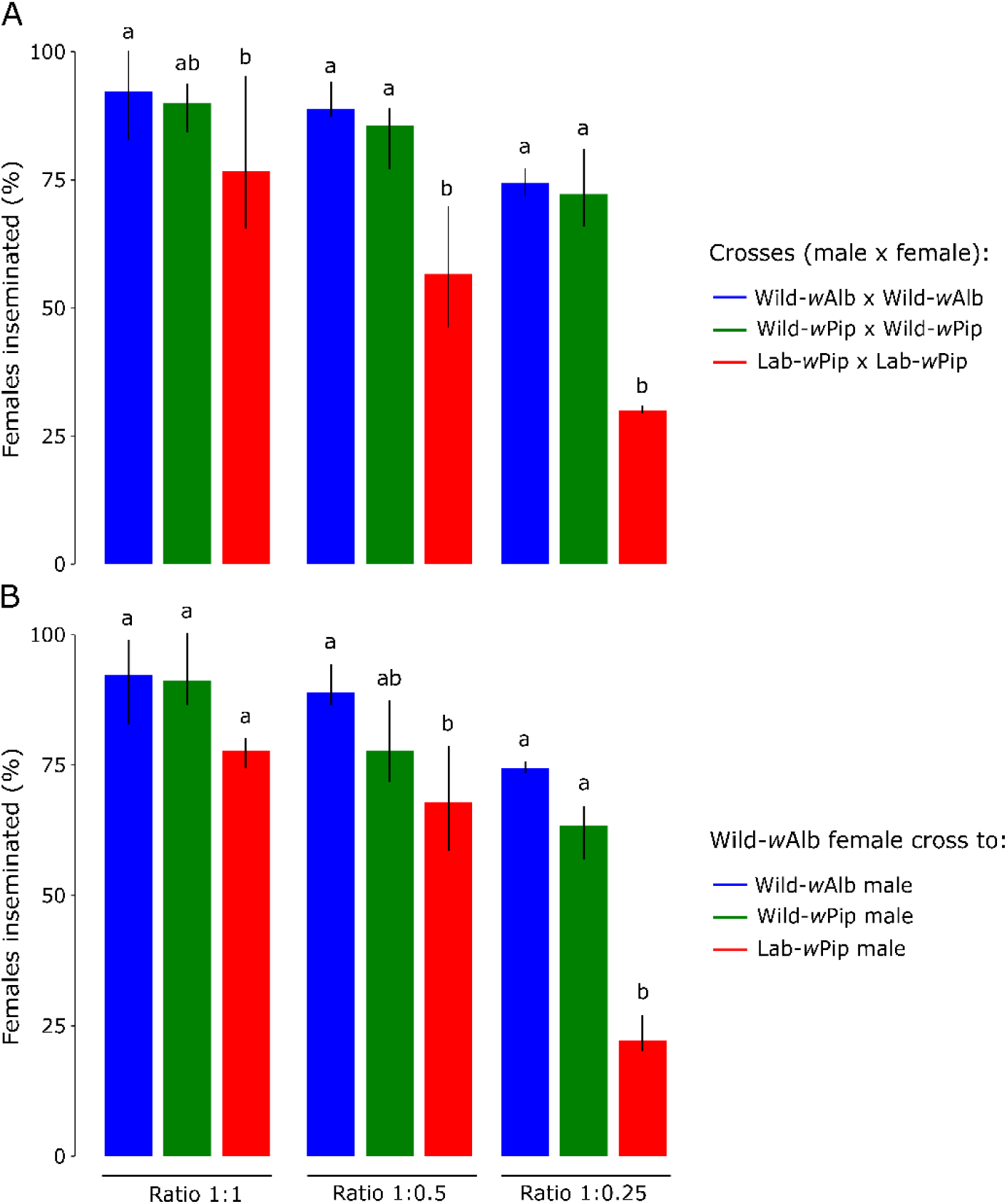
Insemination capacity of incompatible males compared to the wild-*w*Alb males at different male ratio. A: Percentage of inseminated females in compatible crosses (male and female from the same line). B: Percentage of inseminated females in incompatible crosses (male from the lab-*w*Pip and wild-*w*Pip crossed with wild-*w*Alb female). Distinct letters indicate significant variations between the different crosses. The bar plots represent the means, and the vertical black bars the min and max values.

### Mating competitiveness of incompatible males

We compared the mating competitiveness of lab-*w*Pip and wild-*w*Pip lines by crossing virgin wild-*w*Alb females with different ratios of incompatible (from lab-*w*Pip or wild-*w*Pip lines) versus compatible (from wild-*w*Alb line) males. Observed hatch rates in 1 : 0 and 0 : 1 ratios (first and second number corresponding to the ratio of compatible and incompatible males, respectively) were compared with the expected hatch rates under the assumption of a similar mating competitiveness between incompatible and compatible males. We showed that incompatible males from lab-*w*Pip and wild-*w*Pip lines induced a complete CI in crosses with wild-*w*Alb females (ratio 0 : 1 in which only incompatible males were present). Accordingly, for both trans-infected lines, we observed a hatch rate decrease with increasing proportion of incompatible males (Fig. 3). For each ratio, the expected and observed hatch rates in crosses involving wild-*w*Pip males were very close, indicating that these males are as competitive as wild-*w*Alb males. By contrast, lab-*w*Pip males appeared significantly less competitive than wild-*w*Alb males in 1 : 1, 1 : 5 and 1 : 10 ratios (Exact binomial test, *P* < 0.01). Altogether, these data suggest that introgression of the original lab-*w*Pip line results in an increase in male mating competitiveness to levels that are not distinguishable from that of wild-*w*Alb males.

**FIG 3.**
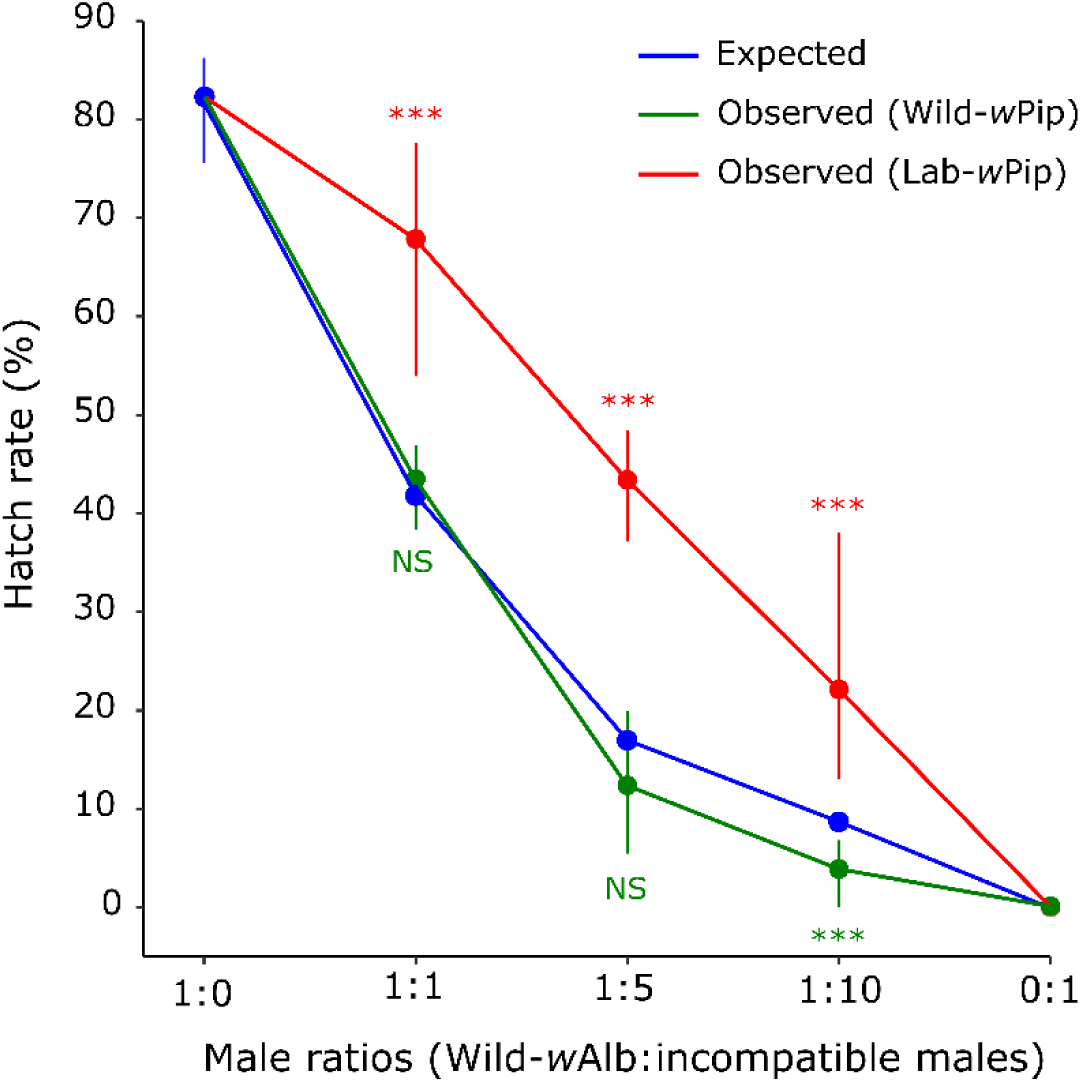
Mating competitiveness of incompatible males from lab-*w*Pip and wild-*w*Pip lines. The blue line indicates the expected hatch rate under a similar competitiveness between compatible and incompatible males. The red and green lines indicate the observed hatch rate with the min and max values. ***P<0.001.

### Evaluation of incompatible males to suppress an Ae. albopictus population

We compared the capacity of lab-*w*Pip and wild-*w*Pip males to suppress an *Ae*. *albopictus* population under laboratory-controlled conditions. In cages where wild-*w*Pip males were introduced weekly, we obtained a suppression efficiency of 100% fifteen weeks after the first release, with a suppression level exceeding 99% from the seventh week after the first release. By contrast, only ∼75% suppression was measured in cages where males from the lab-*w*Pip line were used under the same conditions (Fig. 4A). Similarly, while hatch rates dropped to 0% in cages where wild-*w*Pip males were released, hatch rates plateaued at around 60% in lab-*w*Pip cages and at 80% in control cages (Fig. 4B). In the control cages, where no incompatible males were introduced, the number of eggs laid per week increased substantially over the first seven weeks before plateauing at roughly 5 000 eggs per week, whereas an average value of 74 eggs was recorded in wild-*w*Pip and 1 845 in lab-*w*Pip treatment cages at the end of the experiment (Fig. 4C).

**FIG 4.**
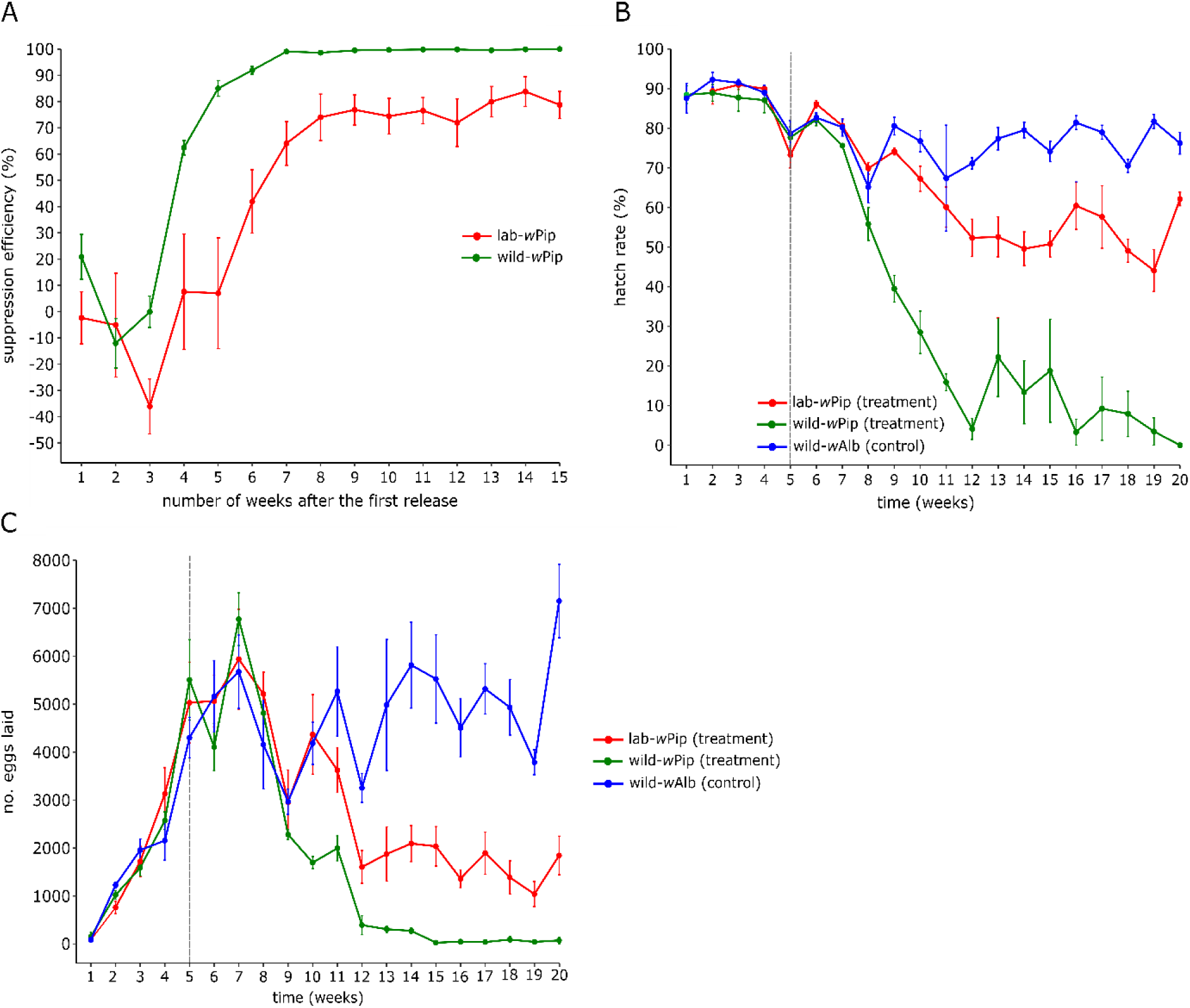
Evaluation of incompatible males from lab-*w*Pip and wild-*w*Pip lines effectiveness to suppress *Ae*. *albopictus* population. A: Suppression efficiency (%). B: Hatch rates of eggs. C: Number of eggs laid. Black vertical dotted lines represent the beginning of incompatible male releases. Each dot represents the mean value, and the vertical lines indicate the standard deviation.

## Discussion

*Wolbachia* infections may alter the fitness of their hosts, with the observed effects influenced by both *Wolbachia* strains and the lineage of the insect host (35, 46, 47). This is particularly true when *Wolbachia* is introduced experimentally into insect hosts, resulting in diverse effects on fitness, which are hardly predictable since densities and tissue distributions can change dramatically upon trans-infection (12, 48). We previously introduced the *w*Pip-IV *Wolbachia* strain in an *Ae*. *albopictus Wolbachia*-free (asymbiotic) laboratory line and showed that this new infection induced a physiological cost, characterized by a decrease in hatch rate and fecundity as compared to a naturally *w*AlbA/B bi-infected or an asymbiotic line (20). The horizontal transfer of *w*Pip into an asymbiotic *Ae*. *albopictus* line from Italy also reduced fitness several generations post-transfer, with the suggestion that a continued selection could attenuate the negative fitness effects associated with artificial *Wolbachia* infection (39, 40).

In our case, we hypothesized that these fitness costs were due to the genetic drift induced by the successive population bottlenecks resulting from the construction of the trans-infected line, including the generation of the asymbiotic line through antibiotic treatment and the selection of isofemale lines showing perfect maternal transmission of *w*Pip (20). Under such circumstances, inbreeding depression is likely to occur through mating between closely related individuals, resulting in increased homozygosity and the appearance of traits associated with reduced fitness (49, 50). Other factors such as gut microbiota could also explain variation of fitness between laboratory and field populations because the microbiome, which can greatly influence *Aedes* mosquitoes LHTs, is drastically affected by laboratory maintenance (51). Indeed, gut microbiota are much less diverse in colonized mosquitoes and tend to be similar in laboratory populations of *Ae*. *aegypti* regardless of geographic origin (52), suggesting that in artificial conditions the nuclear compartment plays a greater role in mosquito performance. For example, in *Ae*. *aegypti*, a loss of fitness due to inbreeding was strongly correlated with a decreased effective population size (53).

In this study we demonstrated that introgression with a wild background lineage for as little as four consecutive generations restored most fitness characteristics of a trans-infected line to those of its native wild counterpart. Indeed, we showed that the post-introgression wild-*w*Pip and the native wild-*w*Alb lines had both comparable fecundity and egg hatch rates. Males from the wild-*w*Pip and the wild-*w*Alb lines also had undistinguishable longevity and insemination capacities, leading to equal mating competitiveness. Finally, we measured the capacity of incompatible males, whether introgressed or not, to suppress a wild population under laboratory-controlled conditions. Using a protocol based on a closed population (no migration) and without taking into account potential density-dependent effects, a population extinction was expected with incompatible males inducing complete CI. The releases of wild-*w*Pip males allowed us to achieve the expected elimination, with complete suppression obtained 15 weeks after the first release, whereas the use of males from the non-introgressed lab-*w*Pip only resulted in suppression levels stagnating at ∼75% until 15 weeks after the first release. An intermediate level of suppression was also obtained in similar conditions when this lab-*w*Pip line was combined with a Genetic Sexing Strain (20). In another investigation, repetitive releases of incompatible males from the triple-*Wolbachia*-*Ae*. *albopictus* HC line were seen to lead to a suppression level close to 100% ∼10 weeks after the first release (15). A limitation of our experiments is that we assessed the effects of introgression only under laboratory conditions. The restoration of incompatible male performance in these conditions does not necessarily indicate similar competitive behavior in the field. Therefore, population suppression experiments should be repeated in semi-field and field conditions before considering the use of this line for a large-scale deployment.

Our findings suggest that several rounds of introgression can be an alternative to a selection process for decreasing the negative impacts of a *Wolbachia* strain post-introduction. It has been shown, for example, in naturally *Wolbachia*-free *Ae. aegypti* as well as in a *w*MelPop-CLA trans-infected line that outcrossing can introduce considerable genetic heterogeneity resulting in improved host fitness (53, 54). Our study also reveals that genetic heterogeneity within a mosquito line, regardless of its geographical origin, is crucial for better characterizing a *Wolbachia*-host association. This is of direct importance when the horizontal transfer aims to select a *Wolbachia* candidate for IIT applications.

Furthermore, to maximize the success of a release program, a common precaution is to introgress the genetic background of wild-collected mosquitoes into the strain to be released through repeated crossing (55–57). This recommendation is also relevant for large-scale production, in which lineage maintenance over multiple generations under laboratory conditions can significantly reduce the fitness and heterozygosity of mosquitoes (53). For example, it has been reported that laboratory-adapted *Aedes* mosquitoes show shorter development and increased body size (53, 58), higher sensitivity to stress (59), and reduced female fecundity relative to their field counterparts (60).

Our findings are highly significant for the implementation of IIT programs as they contribute to improve their efficiency. Introgression should be considered a crucial component in the development of transinfected lines before initiating male release programs. Moreover, in our case, introgression positively impacted female fitness, increasing the number of eggs laid per female, thereby facilitating the mass production process. Because regular introgression by backcrossing during a mass production will be practically challenging, we suggest that introgression should be performed at a laboratory scale before mass production, and that genetic variability could be preserved by the maintenance of a large number of individuals. It has been demonstrated that repeated outcrossing with a wild-type line and maintaining a large effective population size allowed the mass rearing of the triple-*Wolbachia Ae*. *albopictus* HC line up to 15 million mosquitoes per week for over 50 generations without a reduction in the quality of the artificially bred mosquitoes (61).

The data presented in this study are of general interest for release programs, especially for IIT deployment. We demonstrated the potential of introgression with a wild genetic background into a transinfected laboratory line, resulting in the restoration of male fitness that had been artificially infected with *Wolbachia*. The restoration of fitness and genetic variability through introgression is an important development towards the successful application of population suppression programs using this trans-infected line.

## Materials and methods

### Mosquito lines and introgression of a wild genetic background into the trans-infected laboratory line

Three different *Ae*. *albopictus* lines were used in the study. The first one, named lab-*w*Pip, is an *Ae*. *albopictus* line trans-infected with *w*Pip-IV (20), a *Wolbachia* strain naturally occurring in *Culex pipiens* (62). This trans-infected line was originally obtained using a natural population from Reunion Island maintained in the insectary for 40 generations, that was deprived of its natural *Wolbachia* through an antibiotic treatment during six consecutive generations and finally used as a recipient strain of *w*Pip-IV (20). The second line, named wild-*w*Alb and which provided the nuclear background used for the introgression of the incompatible lab-*w*Pip line, was obtained at the city of Le Port, Reunion island, following a single time point sampling using ∼60 ovitraps placed throughout an area of 12.5 hectares. Then, we backcrossed 15 day-old wild-*w*Alb males (using the F_1_ post sampling) with 4-5 day-old females from the lab-*w*Pip line for four consecutive generations, resulting in a line named wild-*w*Pip (expected to display 93,75% of the wild genetic background). Adults of all lines were maintained at a temperature of 27 ±1◦C, a relative humidity of 70 ± 5% and a 12 : 12 h light:dark photoperiod. Females were blood-fed with bovine blood provided by the regional slaughterhouse (Saint-Pierre, La Réunion), supplemented with EDTA (0,1%) and using the Hemotek system (Hemotek Ltd, United Kingdom).

### Verification of CI penetrance of wPip-IV in a wild genetic background

To confirm CI expression following the introgression of the wild-*w*Pip line, we performed reciprocal *en masse* crosses using wild-*w*Pip and wild-*w*Alb lines. Although *en masse* crosses can mask individual variability, especially in the case of moderate CI, we expected complete or nearly complete CI (20), which led us to favor *en masse* rather than individual crosses in order to screen a larger number of mosquitoes. Crosses involved 2-5-day-old virgin females and males (100 from each sex) in 15 × 15 × 15 cm cages (Bugdorm, Taiwan). Three replicates were performed for each crossing experiment. Females were given a blood meal 48h after mating and eggs were collected 5 days later. After 7 days of drying, eggs were allowed to hatch for 48 h (in a jar of 250 mL with tap water supplemented with 5 mg of TetraMin(TETRA)) and the number of hatched and unhatched eggs was counted. A hatch rate was calculated as follows: hatch rate = (number of hatched eggs/total number of eggs) × 100. To account for embryonic mortality not related to CI, we used a corrected index of CI (CI_corr_) (47, 63) calculated as follows: CI_corr_ = [(CI_obs_ − CCM)/(100 − CCM)] × 100, where CI_obs_ is the percentage of unhatched eggs observed in a given incompatible cross, and CCM is the mean mortality observed in the control crosses. The F_2_ and F_5_ were used for wild-*w*Pip and wild-*w*Alb lines, respectively.

### Comparison of life history traits

The larvae-rearing conditions were standardized between lines and was the same for all experiments presented in this study. Specifically, eggs from each line were allowed to hatch for 24 h at 31°C in containers containing 250 ml of water supplemented with 50 mg TetraMin (TETRA). For each line, 2 500 L1 were manually counted and transferred to a tray (53 × 325 × 65 cm, MORI 2A) and fed with a controlled quantity of food (TetraMin (TETRA), day 1: 0.45 g, day 2: 1 g, day 3: 1.25 g, day 4: 1 g, day 5: 1 g, day 6: 1 g) until pupal stage. Male and female pupae were initially separated using a pupae sex sorter (Wolbaki, WBK-P0001-V1 model), and then individually inspected under a binocular loupe.

For each line, longevity was measured by introducing 100 newly emerged males or females separately in 15 × 15 × 15 cm cages (Bugdorm, Taiwan) in which sugar meal (5%) was changed weekly. Three replicates were performed for each line and sex and longevity was determined by recording the number of dead mosquitoes for 45 days.

For each line, the number of laid eggs was measured by placing 200 male and 200 female pupae (one cage per line) in 30 × 30 × 30 cm cages (Bugdorm, Taiwan) and left for 3 days following emergence. A blood meal was then provided and engorged females were placed in a separate cage. Five days later, 50 females were randomly selected and placed individually in a small plastic cup for egg laying. Cups with at least one laid egg were conserved for the analyses. After 7 days of drying, eggs were counted, allowed to hatch for 48 h and hatch rates were measured. The F_3_ and F_6_ were used for wild-*w*Pip and wild-*w*Alb lines, respectively.

The insemination capacity of males was evaluated for each line by placing fifty 2-3-day-old virgin males and females in 15 × 15 × 15 cm cages (Bugdorm, Taiwan). The insemination capacity of males was evaluated by crossing males and females of the same line and crossing incompatible males from the wild-*w*Pip and lab-*w*Pip with the wild-*w*Alb females. Three different sex-ratios: 1:1 (female:male), 1:0.5 and 1:0.25 were tested for each cross. After 48h of mating, all individuals were stored at -20°C. For each cross and sex-ratio, 30 females were randomly selected and dissected to quantify the insemination rate. Females were considered inseminated when presenting at least one spermatheca filled with sperm (64). This experiment was repeated three times, thus 90 females were dissected for each cross and sex-ratio. The F_4_ and F_7_ were used for wild-*w*Pip and wild-*w*Alb lines, respectively.

### Comparison of the mating competitiveness of introgressed vs. non-introgressed males

We evaluated the mating competitiveness of incompatible males by mixing virgin females from the wild-*w*Alb line with different ratios of males from the wild-*w*Pip and lab-*w*Pip lines. Pupae from both lines were allowed to emerge in separate 30 × 30 × 30 cm cages (Bugdorm, Taiwan) with sugar meal (5%); 2-3-day old virgin females and males were used for this experiment. Females (N = 100) were first placed inside cages followed by the simultaneous release of all males, and mating was allowed for 48 h. All males were then removed, a blood meal was provided and eggs were collected by oviposition *en masse* 5 days later. After 7 days of drying, eggs were allowed to hatch and the hatching rate was measured and used as a proxy of mating competitiveness. Five ratios were tested: 1 : 1 (50♂ wild-*w*Alb : 50♂ wild-*w*Pip or lab-*w*Pip), 1 : 5 (17♂ wild-*w*Alb : 85♂ wild-*w*Pip or lab-*w*Pip), 1 : 10 (9♂ wild-*w*Alb : 90♂ wild-*w*Pip or lab-*w*Pip) and two control ratios, 1 : 0 (100♂ wild-*w*Alb : 0♂ wild-*w*Pip or lab-*w*Pip) and 0 : 1 (0♂ wild-*w*Alb : 100♂ wild-*w*Pip or lab-*w*Pip). Three replicates were performed for each cross and ratio. The F_3_ and F_6_ were used for wild-*w*Pip and wild-*w*Alb lines, respectively.

### Laboratory cage suppression experiment

A wild population was established in each control cage (4 replicates) introducing weekly 100 males and females pupae of the wild-*w*Alb line from the beginning to the end of the experiment. The treatment cages (4 replicates) were treated similarly until the end of week 4, then 500 incompatible adult males from wild-*w*Pip or lab-*w*Pip line were released weekly, resulting in a 5 : 1 ratio (500 wild-*w*Pip or lab-*w*Pip males : 100 wild-*w*Alb males). We took into account the level of population suppression induced by the repeated releases of incompatible males in the treatment cages. Thus the number of introduced wild-*w*Alb pupae was determined by the difference of egg hatch rates between control and treatment cages. For example, if egg hatch rate was 80% and 60% (20% difference) in control and treatment cages, respectively, we introduced 20% less wild-*w*Alb males and females in the treatment cages the following week (100 × 0.2 = 80), therefore 80 male and female pupae, instead of 100 pupae of each sex in the control cages.

Throughout the experiment, 3 blood meals were provided weekly and eggs were collected once a week. After 7 days of drying, all eggs were counted and allowed to hatch for 24 h. Thus, the number of eggs and hatch rates were determined weekly for each cage. Altogether, 12 cages (45 × 45 × 45 cm, Bugdorm, Taiwan) were monitored, including 4 repetitions for each cage type (control, wild-*w*Pip and lab-*w*Pip). The suppression efficiency (SE) was calculated based on the following equation, where Hc and Ht are the numbers of hatched eggs counted weekly in control and treatment cages respectively (15, 19). SE (%) = [(µHc – Ht) / µHc] x 100. The F_4_ and F_7_ were used at the beginning of the experiment, for wild-*w*Pip and wild-*w*Alb lines, respectively.

### Statistical analyses

Longevity data were analysed using a log-rank test. Fecundity data (count data) were analysed using a GLMM (quasi-poisson family, log link) in which the different mosquito lines were included as a fixed effect and the repetitions (individual egg laying) as a random factor. We used a generalized linear mixed model (GLMM) to analyse egg hatch rate (binary data, quasi-binomial family, logit link) and insemination rate data (binary data, binomial family, logit link). The overdispersion of the data was checked using a R code proposed by Ben Bolker and others (https://bbolker.github.io/mixedmodels-misc/glmmFAQ.html). A F-test for the GLMM-quasi-binomial and quasi-poisson models were used to analyse deviance. Exact binomial tests were used to compare the observed and expected hatch rates in the mating competitiveness experiment (proportional data). Analyses were performed in R version 4.3.3 (65), using lme4 package for all mixed models (66), MASS package (67) for using the quasi-binomial and quasi-poisson families in the GLMMs, and survival package for longevity data analyses (68). For all data, the significance level was set to α = 0.05.

## Acknowledgements

We thank Dr. David Wilkinson for his helpful proofreading of the manuscript.

## Funding

This work was supported by a Credit Impôt Recherche provided by the French Ministry of research and higher education to SymbioTIC and by a Fonds européen de développement régional (FEDER) (INTERREG VI program; OPERATING SEYWOL n°: REU004954).

